# Optimisation of TP53 reporters by systematic dissection of synthetic TP53 response elements

**DOI:** 10.1101/2023.07.24.549988

**Authors:** Max Trauernicht, Chaitanya Rastogi, Stefano G. Manzo, Harmen J. Bussemaker, Bas van Steensel

## Abstract

TP53 is a transcription factor that controls multiple cellular processes, including cell cycle arrest, DNA repair, and apoptosis. The relation between TP53 binding site architecture and transcriptional output is still not fully understood. Here, we systematically examined in three different cell lines the effects of binding site affinity and copy number on TP53-dependent transcriptional output, and also probed the impact of spacer length and sequence between adjacent binding sites, and of core promoter identity. Paradoxically, we found that high-affinity TP53 binding sites are less potent than medium-affinity sites. TP53 achieves supra-additive transcriptional activation through optimally spaced adjacent binding sites, suggesting a cooperative mechanism. Optimally spaced adjacent binding sites have a ∼10-bp periodicity, suggesting a role for spatial orientation along the DNA double helix. We leveraged these insights to construct a log-linear model that explains activity from sequence features, and to identify new highly active and sensitive TP53 reporters.

## INTRODUCTION

The transcription factor TP53 is a master tumour suppressor that plays a crucial role in the regulation of cell growth and division. Upon induction by diverse stress signals, TP53 becomes activated and binds to DNA to regulate the expression of genes involved in cell cycle regulation and DNA repair. Typically, TP53 binds DNA as a tetramer over a 20-bp binding site (BS) consisting of two decameric half-sites (2x RRRCWWGYYY, R = A/G, W = A/T and Y = C/T) with an optional short spacer in between them (1). Since TP53 can bind hundreds of target genes with diverse functions (2), selective target gene regulation is important for correct TP53 function. One working hypothesis is that TP53 achieves selectivity by recognition of specific BS variants (3-6).

Certain TP53-responsive promoters have been used to construct artificial reporter genes that are now widely used to monitor the activity of TP53 in cells (7-11) and in transgenic mice (12). Such reporters have been invaluable for our understanding of the response of TP53 to a variety of stress signals. However, it is unclear whether these reporters may be further optimised to achieve increased sensitivity. More accurate and sensitive detection of TP53 activity would allow measurements of subtle changes in TP53 transcriptional regulation in small cell populations, and potentially identify and classify different TP53 mutations with decreased transcriptional activity (13,14). Generating better reporters requires a better understanding of the logic of promoter regulation by TP53 BSs.

Genomic TP53 BSs show a highly varying degree of deviation from the optimal sequence (15,16). Therefore, several studies have characterised the sequence requirements for TP53 function in order to establish the relationship between TP53 BS architecture and transcriptional activity. A study based on TP53 reporter assays established that genomic TP53 BSs can have up to 1,000-fold differences in transactivation capacity and that genomic response elements containing two BSs are more active than those with a single BS (17). It was also shown that TP53 BSs without a spacer between the decameric half-sites lead to more potent transcriptional activation (9), and that the spacer length between two adjacent TP53 BSs can influence transcriptional output (9,18). Additionally, it was reported that TP53 transcriptional activation correlates positively with TP53 binding affinity at high TP53 levels, whereas transactivation at low TP53 levels can be explained by the torsional flexibility of the TP53 BS (13,14). More precisely, TP53 BSs with a ‘CATG’ sequence at the center of their decameric half-sites were shown to be more torsionally flexible, allowing for stable binding of TP53 dimers (19) and leading to strong transcriptional activation even at low levels of TP53 (13).

Recently, two studies systematically probed the transactivation capacities of genomic TP53 BSs using massively parallel reporter assays (MPRAs). In one, it was demonstrated that TP53 drives transcription autonomously (i.e., without requiring co-factors) and that the presence of a single canonical TP53 BS is sufficient to drive strong activation (20). In another MPRA-based study, it was shown that co-factors and the sequence context of the TP53 BS can alter TP53-dependent transcriptional activity substantially (21).

While these functional studies revealed important links between the architecture of TP53 response elements and transcriptional activity, we are still lacking detailed understanding of the grammar underlying TP53-mediated transcriptional activation.

A more detailed understanding of the sequence requirements of TP53-driven transcription may be obtained by designing and probing synthetic sequences that do not exist in the genome. This allows for controlled and isolated evaluation of one BS sequence feature at a time without confounding factors. This approach was taken for several other TFs previously and has proven useful to understand how TFs drive transcription (22-24). We reasoned that this may also uncover rules that can be applied to design more sensitive reporters for the monitoring of TP53 activity.

Here, we took such a fully synthetic approach to evaluate the relation between TP53 BS architecture and transcriptional activity. We systematically designed more than a thousand TP53 reporters and performed MPRAs to probe the effects of TP53 BS affinity, BS copy number, BS spacing, and core promoter-interaction on transcriptional output. We find that all investigated features substantially alter transcriptional output. The strongest effect on transcriptional activity was caused by BS copy number and the spacer length between adjacent BSs, suggesting that TP53 molecules can co-operatively drive transcription from adjacent BSs, if positioned correctly. Together, our results provide new insights into how TP53 drives transcription. We leverage this knowledge to design optimised TP53 reporters with substantially higher sensitivity than a commonly used reporters.

## MATERIALS & METHODS

### TP53 reporter library design

To select TP53 BSs with a large range of binding affinities, we used a pre-built full-length TP53 *No Read Left Behind* model (25) with a 24-bp footprint that includes 2-bp up- and down-stream of the canonical 20-bp TP53 BS. The binding energy coefficients (ΔΔG/RT) underlying this model were previously found to be highly consistent (R^2^ = 0.84) with those independently inferred from data for a C-terminal domain (CTD) truncation of TP53, providing confidence in its quantitative predictive value. We used it to generate four synthetic BSs with minimal edits and model-predicted relative affinities ranging from 6% (BS006) to 100% (BS100) using a simple sequence sampling algorithm. Additionally, a zero-affinity BS was generated by mutating the C and G bases in the CWWG cores of both half-sites. BS sequences are shown in **Figure 1A**. Four copies of the resulting five BSs were then placed upstream of the core promoter. Also, BSs were combined with BSs that have one affinity higher or lower (e.g., BS100 with BS037 or BS006 with BS000) in all possible combinatorial arrangements across the four positions (e.g., BS100-BS037-BS037-BS100-core promoter or BS037-BS100-BS037-BS100-core promoter).

**Figure 1:**
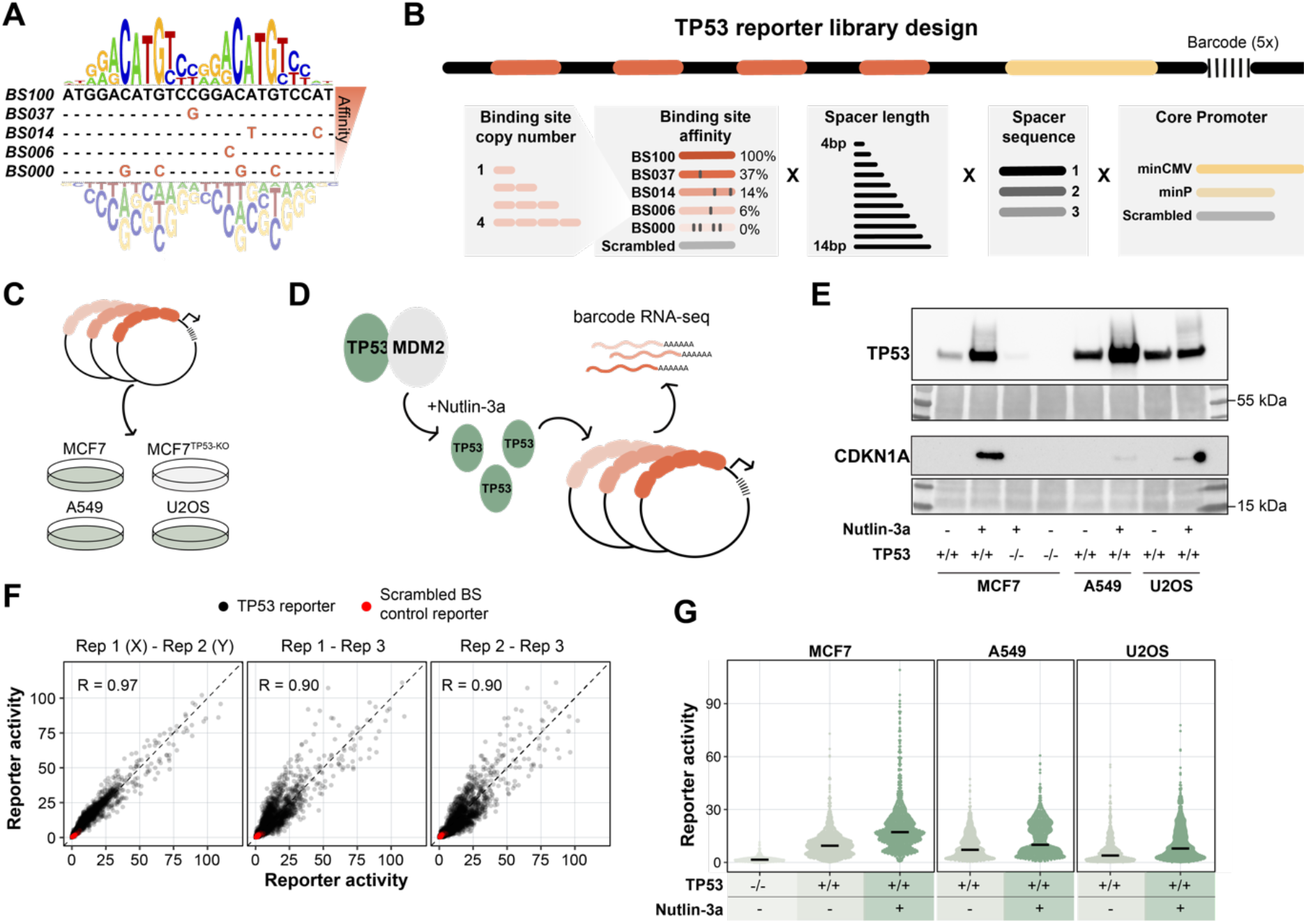
Systematic screening of TP53 reporter variants. (**A**) The 24-bp position weight matrix (PWM) underlying the TP53 affinity model (25), which was used to design five TP53 BSs with a broad range of predicted binding affinities. (**B**) Scheme of the TP53 reporter design. TP53 BSs were placed at four positions, with the first BS positioned 10 bp upstream of the core promoter. TP53 BSs, BS-BS spacer length, BS-BS spacer sequence, and core promoter were varied as indicated in the scheme. (**C**) The TP53 reporter plasmid library was transfected into A549, U2OS, and TP53 proficient or TP53-KO MCF7 cells. (**D**) Upon transfection the cells were treated with Nutlin-3a or vehicle control. (**E**) Western blot showing the protein levels of TP53 and CDKN1A in all tested conditions. Bottom panels show Ponceau S staining serving as loading control. (**F**) Pairwise correlations between the three independent replicates of the computed reporter activities. Reporter activities were computed from the barcode counts in the mRNA, and are displayed as fold enrichment over the mean of the scrambled BS controls per core promoter and condition. Only data for MCF7 cells are shown. Experiments in A549 and U2OS cells were performed in two independent biological replicates which correlated highly (Pearson’s R = 0.99 and 0.98 for A549 and U2OS, respectively). (**G**) Reporter activities (average of the three independent replicates) of all TP53 reporters compared to the scrambled BS control reporters in the tested conditions. Black horizontal lines indicate median values.

Then, three different spacer sequences with lengths ranging from 0-10 bp were designed. 2,000 random sequences with a GC content of 40-60% were generated (sim.DNAseq function in R from package SimRAD (version 0.96)), these sequences were then scanned across all spacer lengths in combination with 4 bp of the left and right side of the five TP53 BSs using FIMO (26). Nine sequences without any significant TF binding (p-value threshold 1e-4) were selected and placed in between the TP53 BSs (three different spacer sequences per reporter, times the three spacer sequences). A similar approach was taken to generate three 10-bp spacer sequences in front of the core promoter.

The two core promoter sequences, minCMV (27) and minP (derived from pGL4 (Promega, Madison, WI, USA)), were placed downstream of the 10-bp spacer sequence, followed by a S1 Illumina adapter sequence and a unique 13-bp random barcode sequence (each unique construct was linked to five different barcodes). All generated random barcodes had a Levenshtein distance of at least three with respect to one another and barcodes with an unbalanced GC ratio were removed (create.dnabarcodes function from the R package DNABarcodes (version 1.2.2) (28)).

To include TP53-independent background controls, a scrambled TP53 BS was generated while excluding unwanted TF binding using FIMO, and placed in all four BS positions in front of the core promoter. Similarly, a scrambled 32-bp core promoter sequence was included as control.

To be able to compare the transcriptional levels of the synthetic TP53 reporters to commonly used TP53 reporters, we included a frequently used TP53 response element as a benchmark control. This TP53 response element is controlling the *GADD45A* gene (7) and is commercially available (pGL4.38, Promega). The benchmark response element was also linked to the three selected spacer sequences and the two core promoters. All reporter sequences were completed with 18-bp primer adapter sequences (that were also scanned using FIMO) in both flanks for cloning purposes. The resulting sequence pool had a total length of on average 237 bp (at least 205 bp up to 269 bp) and was ordered as oligonucleotide library from Twist Biosciences.

### Cloning of the TP53 reporter plasmid library

First, the entry vector was constructed by adding a bcl-2 splice acceptor and four SV40 polyA-signals to a plasmid backbone based on pSMART (Addgene #49157) containing a green fluorescent protein (GFP) open reading frame followed by a SV40 derived polyadenylation signal (PAS) (29). This step is done to reduce background signals deriving from the ORI (30). The oligonucleotide library was resuspended in TE buffer (Invitrogen) to a final concentration of 20 ng/μl. 10 ng of the oligonucleotide library was then PCR amplified (1’ 95°C, 6x(15’’ 95°C, 15’’ 57°C, 15’’ 72°C), 1’ 72°C) by MyTaq Red mix (Bioline) using primers that add overhangs with EcoRI (MT024, **Supplementary Table S1**) or NheI (MT025) restriction enzyme sites. The PCR product was then purified using CleanPCR (CleanNA) beads at 1.8:1 beads:sample ratio, then digested with EcoRI-HF (NEB, #R3101) and NheI-HF (NEB, #3131) by incubating the PCR product at 37°C for 1 h, and then again bead purified as before. 1 μg of the entry vector was also digested with EcoRI-HF and NheI-HF and the linearised product was purified from a 2% agarose gel using PCR Isolate II PCR and Gel Kit (Bioline). The digested and purified reporter pool was then ligated into 80 ng of the linearised entry vector using Takara ligation kit v1.0 (#6021; Takara) at a 1:3 (vector:insert) ratio. The ligation mix was then bead purified as before and transformed into MegaX DH10B T1R Electrocomp™ Cells (Invitrogen) using 1 μl of the ligation mix. The library complexity was estimated from plated serial dilutions of the transformed cells to be ∼300,000 colony forming units. Transformed cells were transferred to 200 ml standard Luria Broth (LB) plus kanamycin (50μg/ml), grown overnight and purified using a Maxi plasmid purification kit (#12162; Qiagen).

### Reporter library transfection and cell culture

MCF7 cells (ATCC HTB-22) and a polyclonal line of MCF7-TP53-KO cells were a kind gift from Karin de Visser (Netherlands Cancer Institute). MCF7-TP53-KO was generated using a sgRNA targeting exon 4, yielding >90 % of +1 insertions (31). A549 cells (ATCC CCL-185), MCF7 cells and MCF7-TP53-KO cells were cultured in DMEM medium (GIBCO) supplemented with 10% fetal bovine serum (FBS, Sigma). U2OS (ATCC HTB-96) cells were cultured in McCoy’s 5a Medium (Gibco) supplemented with 10% FBS (Sigma). All cells were routinely tested for mycoplasm. Per transfection, 1×10^6^ cells were resuspended in transfection buffer (100 mM KH2PO4, 15 mM NaHCO3, 12 mM MgCl2, 8 mM ATP, 2 mM glucose (pH 7.4)) supplied with 1 μg of plasmid library and electroporated in an Amaxa 2D Nucleofector using program E-014. After nucleofection, cells were resuspended in 2 ml complete medium and plated in 6-well plates. For stimulation with Nutlin-3a, 8 μM of Nutlin-3a (MedChemExpress, #HY-10029) was added directly after transfection. 24 hours after transfection, cells were harvested and resuspended in 800 μl TRIsure (#BIO-38032; Bioline) and stored at -80 °C until further use. Transfections were done in three independent replicates on three separate days for MCF7 and MCF7-TP53-KO cells and in two independent replicates on two separate days for U2OS and A549 cells.

### RNA extraction, reverse transcription and barcode amplification

RNA extraction was done using the standard procedure according to the TRIsure protocol. After RNA extraction, 2 μg of RNA was treated with DNase I for 30 minutes (#04716728001; Roche) and subsequently treated with 1 μl 25 mM EDTA at 70 °C for 10 minutes to inactivate DNase I. cDNA was synthesised by addition of 1 μl gene-specific primer targeting the GFP ORF (10 μM, MT165) and 1 μl dNTPs (10mM each) followed by incubation at 65 °C for 5 minutes. Then, the reverse transcription reaction was set up by adding 4 μl of RT buffer, 20 units RiboLock RNase inhibitor (#EO0381; ThermoFisher Scientific), 200 units of Maxima reverse transcriptase (#EP0743; ThermoFisher Scientific) and 2.5 μl of water. The reaction was then incubated for 30 minutes at 50 °C followed by heat-inactivation at 85 °C for 5 minutes. 20 μl of cDNA were then PCR amplified (1′ 96 °C, 20x(15″ 96 °C, 15″ 60 °C, 15″ 72 °C)) in a 100 μl reaction using MyTaq Red mix and primers containing the Illumina S1 and p5 adapter (MT397) and the Illumina S2 and p7 adapter (MT164). To generate input plasmid DNA (pDNA) barcode counts that serve as normalisation control, the plasmid library that was used for the transfections was linearised using EcoRI-HF and subsequently 1 ng of linearised vector was PCR amplified as before using 8 cycles. PCR products were pooled and purified by double-sided bead purification using beads:sample ratios of 0.6:1 followed by 1.2:1 on the supernatant. The sequencing library was then sequenced using a 75 bp single-read NextSeq High Output kit (Illumina).

### TP53 and CKDN1A Western blotting

70% confluent 10 cm dishes of A549, U2OS, MCF7 or MCF7-TP53-KO cells were treated for 24 hours with 8 μM Nutlin-3a or DMSO before harvesting and resuspending in 100 μl protein isolation buffer (50 uL HNTG buffer (50 mM HEPES pH 7.4, 150 mM NaCl, 0.1% Triton X-100, 10% Glycerol), 50 uL RIPA buffer (50 mM Tris-HCl pH 7.4, 150 mM NaCl, 1mM NaF, 1% sodium deoxycholate, 1% Triton X-100, 0.1% SDS), supplemented with cOmplete Protease Inhibitor Cocktail (Roche)). Resuspended cells were then incubated for 30 minutes at 4 °C in a spinning wheel, before centrifugation for 20 minutes at 12,000 rpm at 4 °C. The supernatant was kept at -80 °C until further use. Protein concentration was determined using Pierce™ BCA Protein Assay Kit (Invitrogen). 50 μg of total protein per condition were used as input for Western blotting. Western blotting was performed according to standard procedures using the following antibodies and dilutions: TP53 (DO-1, 1:500, Cell Signaling #18032, mouse), CDKN1A (F-5, 1:1000, Santa Cruz Biotechnology, #sc-6246, mouse).

### Reporter activity computation and normalisations

Raw barcode counts were clustered using *starcode* (32) using a Levenshtein distance of 1. Next, clustered barcode counts were normalised by library size. To be more precise, a pseudocount of 1 was added to the barcode counts before dividing this number by the total sum of all barcode counts per sample per million. From these normalised barcode counts activities were computed by dividing the cDNA barcode counts by the pDNA barcode counts. The activities were normalised by dividing the activities by the mean of the activities of the scrambled BS reporters per core promoter and sample. Normalised activities were then averaged over the five different barcodes and finally over the three independent replicates.

### Log-linear model of reporter activities

To simplify the log-linear model, only reporters with four identical BSs were considered. To accommodate the observed spacing effects, the BS-BS spacer length was transformed from simple bp to a helical positioning score as follows:

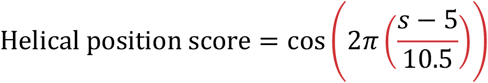

where *s* denotes the BS-BS spacer length in bp. The BS-BS spacer length was subtracted by five to centre the cosine function around a spacer length of 5 bp. After transformation, a log-linear model was fit using the following expression:

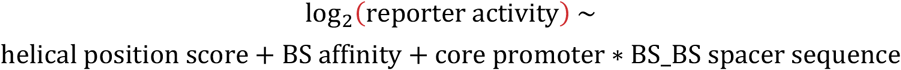

where BS affinity, core promoter, and BS-BS spacer sequence were used as categorical variables. Models were fit using the lm function in R from the stats package (version 3.6.2).

## RESULTS

### A reporter library to dissect TP53 regulatory logic

#### Library design

To investigate the relation between TP53 response element architecture and transcriptional output, we designed a barcoded reporter library consisting of >1,200 unique TP53 reporters. The reporters were systematically designed to capture a broad range of predicted TP53 binding affinities. We selected four TP53 binding sites (BSs) with a predicted relative affinity of 6% (BS006), 14% (BS014), 37% (BS037), or 100% (BS100) of the optimal BS, as estimated by computational modelling of *in vitro* binding assay data (25) (**Figure 1A**). The full 20-bp motif binds a TP53 tetramer (33). We also included a BS with four mutations at the most essential positions, which served as ‘zero-affinity’ control (BS000). We then designed the full TP53 reporter sequences by placing unique combinations of the five BS variants into four different positions on the reporter (**Figure 1B**). More precisely, BSs were only combined with BSs that have one affinity higher or lower (e.g., BS037 with BS014), which also allows to investigate the effect of BS copy number using the BS006-BS000 combinations. The BSs were separated by one of the three distinct BS-BS spacer sequences that were designed to minimise predicted affinity for any known other TF (see Methods). We also varied the spacer length between individual BSs from 4-14 bp to test the relation between BS positioning and transcriptional output. The resulting sequences were then coupled to one of two different core promoters (a minimal TATA promoter (derived from pGL4, Promega) or a minimal CMV promoter (27)). Downstream of the core promoter we placed a ∼2 kb transcription unit containing a unique 13-bp barcode that enabled the detection of all reporter constructs in parallel. Furthermore, we included reporters with scrambled BSs and scrambled core promoters, serving as negative controls. To benchmark our designed reporters, we also included a frequently used and commercially available TP53 reporter sequence (pGL4.38, Promega) (7), that consists of a genomic TP53 response element containing two TP53 BSs. This sequence was coupled to the three spacer sequences and two core promoters used in this study, yielding six benchmark reporters. The resulting library included 1,225 different designs in total. To minimise biases caused by individual barcodes, we used five different random barcodes with each design, the results of which were averaged.

#### Probing multiple cell types and TP53 concentrations

We transiently transfected this library into the three cell types A549 (lung carcinoma), U2OS (osteosarcoma) and MCF7 (breast adenocarcinoma) to probe the reporters across distinct molecular backgrounds and TP53 concentrations. As a negative control, we transfected the library into a polyclonal MCF7 TP53-knockout (TP53-KO) cell line (31) (**Figure 1C**). Additionally, we treated the cells upon transfection with Nutlin-3a (**Figure 1D**). This compound inhibits MDM2, which is a negative regulator of TP53, and thereby stabilises and activates TP53 (34). By Western blotting we confirmed the near-absence of TP53 in the TP53-KO cells and induction of TP53 levels in the presence of Nutlin-3a, the latter leading to activation of the TP53 target gene *CDKN1A* (**Figure 1E**). 24 hours after transfection, we quantified the barcode abundancies in the mRNA by high-throughput sequencing. We next calculated the reporter activity by normalizing the barcode counts in the mRNA to the barcode counts in the input plasmid DNA. The activities were then scaled per sample by calculating the enrichment over the background activity of the scrambled BS control reporters. This normalisation was done separately per core promoter.

#### Overview of data

We performed the reporter assay as three independent biological replicates for MCF7 cells and two independent replicates for A549 and U2OS cells, which all showed high pairwise correlations (Pearson’s R = 0.90-0.99) (**Figure 1F**). We averaged these replicates for further downstream analysis. As expected, we observed a clear correlation between cellular TP53 levels and the reporter activities (**Figure 1G**): the median activity of the reporters carrying at least one functional BS was 6-fold higher in the MCF7 TP53-WT cells compared to the TP53-KO cells, and further increased 1.8-fold upon stimulation with Nutlin-3a. Similarly, activity of TP53 reporters increased 1.4- and 2-fold upon stimulation with Nutlin-3a in A549 and U2OS cells, respectively. Interestingly, in TP53-WT cells the activity of the individual reporters varied more than 50-fold (**Figure 1G**), indicating that the precise design can have dramatic impact on the transcriptional output.

### Paradoxical effect of BS affinity on reporter activity

#### BS affinity does not simply correlate with activity

To identify the determinants of activity, we first analysed the effect of BS affinity on reporter activity by comparing sets of reporters that carry four BSs of the same affinity. Surprisingly, the reporter activity did not show a simple correlation with BS affinity (**Figure 2A**): without Nutlin-3a treatment, optimal activity was not obtained with the highest-affinity BS100, but rather with BS014, which has 7-fold lower affinity. This was observed across all three tested TP53-WT cell types (**Figure 2A**). BS014 also displayed activity above background in MCF7 TP53-KO cells. This is likely due to the presence of a small proportion of TP53-WT cells in the polyclonal TP53-KO population (note the faint band in the TP53 Western blot in the presence of Nutlin-3a, **Figure 1E**). Interestingly, even the lowest-affinity BS006 showed strong activation in MCF7 and A549, but not U2OS. Possibly this can be explained by a lower level of TP53 transcripts (Human Protein Atlas, https://v23.proteinatlas.org/ENSG00000141510-TP53/cell+line) and TP53 proteins (DepMap, https://depmap.org/portal/download/all/, (35)) in U2OS cells. Treatment with Nutlin-3a increased reporter activities across all BSs, with BS014 being the most active in A549, U2OS, and BS006 being the most active in MCF7. These results indicate that increasing the BS affinity does not necessarily lead to higher transcriptional output. Possibly, at higher affinities BSs become saturated at basal TP53 levels, and the activity of the reporters appears to be compromised when occupancy by TP53 becomes too high.

**Figure 2:**
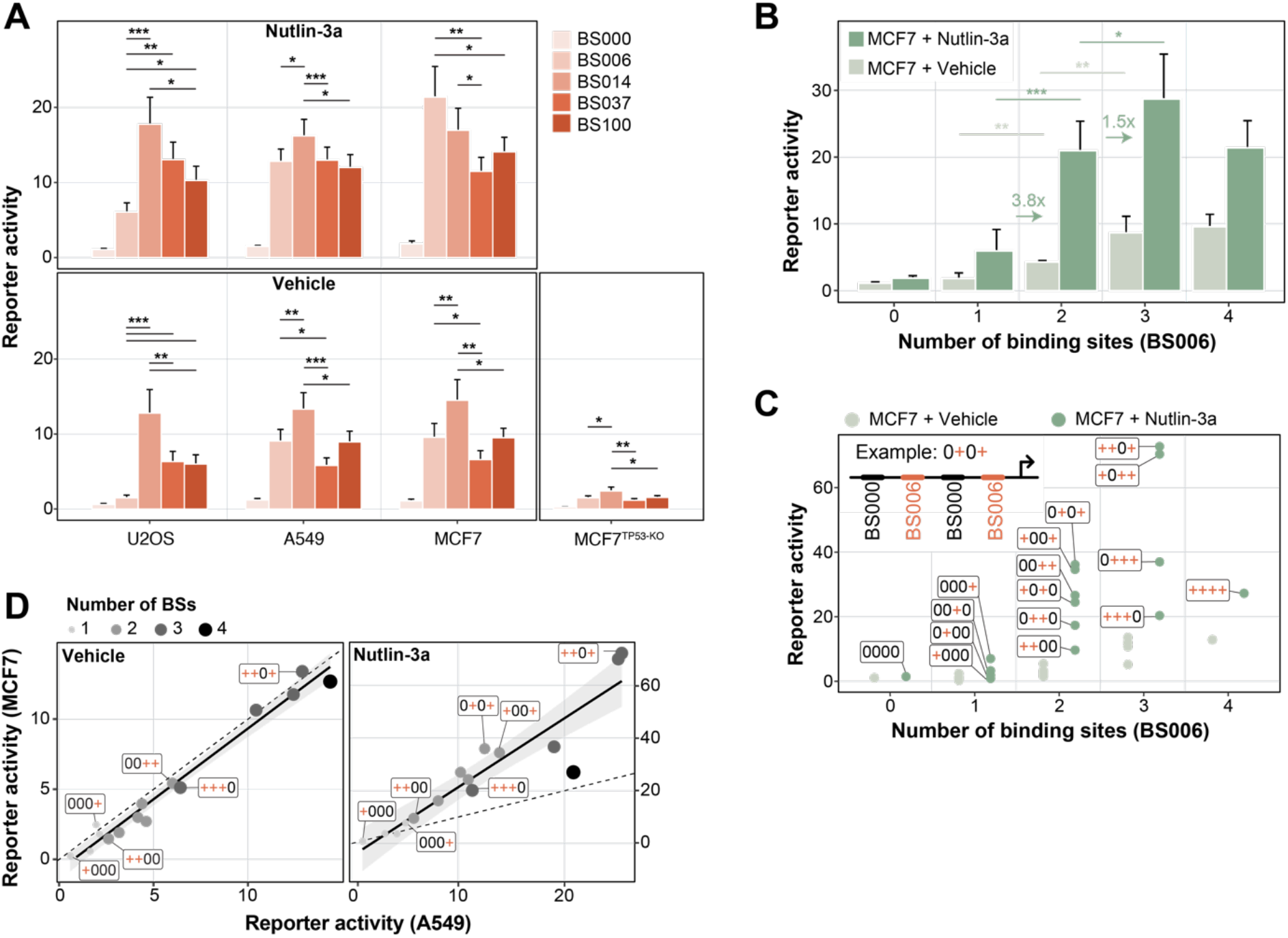
The impact of BS affinity and copy number on reporter activity. (**A**) Reporter activity per condition and BS, as indicated by the colour. Only reporters with four identical copies of the BS and a BS-BS spacer length of 11 bp between all BSs are displayed. Bars indicate the mean and error bars the standard deviation of reporter activities across all BS-BS spacer sequences and core promoters. P-values were calculated using two-tailed paired Student’s t-test: significance levels: ***: p < 0.001, **: p < 0.01, *: p < 0.05. (**B**) Impact of BS006 copy number on reporter activity in TP53-WT cells and TP53-WT cells stimulated with Nutlin-3a. Displayed is the mean and standard deviation of reporter activities across all BS-BS spacer sequences and core promoters. The arrows and numbers indicate the fold-change in reporter activity from one to two or two to three copies of BSs in the MCF7 + Nutlin-3a condition. P-values were calculated using two-tailed paired Student’s t-test: significance levels: ***: p < 0.001, **: p < 0.01, *: p < 0.05. (**C**) Reporter activity as function of copy number and positioning of the BSs. The cartoon depicts the BS code of the reporters; + (red) = BS006, 0 (black) = BS000, arrow = core promoter. Displayed is the mean of the minCMV reporter activities across all BS-BS spacer sequences. (**D**) Correlation of the data plotted in **C** between MCF7 and A549 cells without stimulation (left panel) and with stimulation of Nutlin-3a (right panel).

#### Low affinity of TP53 BS in the human genome

We wondered how the affinities of the tested BSs compare to those of endogenous TP53 BS in the human genome. To address this, we used the same algorithm (25) to calculate affinities of TP53 BSs in a set of previously identified chromatin immunoprecipitation (ChIP) peaks (20). Strikingly, the vast majority of these endogenous BSs has even lower predicted affinities for TP53 than BS006 (**Supplemental Figure 1A**). This is in agreement with earlier observations that endogenous BSs usually do not match the consensus TP53 BS (16). This may indicate that also in the native genomic context low-affinity BSs function better than high-affinity BSs, although it seems likely that cooperative interactions with other TFs alter the effective affinities in this context.

#### Synergy between BSs

Next, we investigated the relationship between the number of TP53 BSs and the transcriptional output. For this we focused on reporters with combinations of BS006 and BS000. We first looked at the effect of adding BSs from the promoter-proximal position. As expected, in MCF7 TP53-WT cells the transcriptional activity increased with the copy number of BS006 (**Figure 2B**). A similar effect was observed in MCF7 cells treated with Nutlin-3a, although the reporter activity was generally higher. Interestingly, in cells stimulated with Nutlin-3a, addition of a second BS yielded on average a 3.8-fold increase relative to reporters with only one BS. Hence, the effect of adding a second BS is supra-additive, suggesting that two TP53 tetramers can synergise to boost transcriptional output. A third BS increased the transcriptional activity further, although not to the same extent, and reporters with four BSs were on average equally active as reporters with two BSs (**Figure 2B**). This dose-response relationship was slightly different in the absence of Nutlin-3a, where the addition of a third BS led to a relatively stronger transcriptional increase (**Figure 2B**). We note that TP53 targets in the human genome rarely have more than two BS (**Supplemental Figure 1B**), and in such instances the spacing of two BSs varies more widely than the spacing range we used in our reporter set (**Supplemental Figure 1C**). Nevertheless, as we will show below, studying arrays of up to four BSs provides useful information for the design of improved reporters.

#### Positioning of BS affects activity

Next, we investigated the complete combinatorial pool of BS000-BS006 reporters to test if the relative positioning of BS006 had an impact on transcriptional output in MCF7 cells. Interestingly, in the presence of Nutlin-3a the activity of reporters with the same copy number of BSs showed up to 3.6-fold variation in activity, depending on where the BSs are positioned (**Figure 2C**). This indicated that not only the copy number, but also the position of the BSs is important. In reporters with only one BS, activity was highest if the BS was positioned close to the core promoter. Similarly, in reporters with two or three BSs, those with a BS immediately next to the promoter were most active. Hence, TP53 appears more potent when the most proximal BS is placed 10 bp from the start of the core promoter instead of 41 bp.

#### Excessive BS density is counter-productive

The highest activity was obtained with three rather than four BSs, but only when the array of three BSs was interrupted by a zero-affinity BS000 (i.e., ++-+ or +-++ in **Figure 2C**), which effectively increases the spacing between two functional BS from 11 to 42 bp. These findings highly correlated with the positional effects observed in A549 cells (**Figure 2D**), although for the latter the dynamic range in the Nutlin-3a condition was reduced. Together, these results highlight the importance of BS positioning and suggest that too dense occupancy of TP53 can lead to reduced transcription activity.

### Effects of BS spacing

#### BS-BS spacer length impacts transcriptional output

To examine the effect of BS spacing further, we analysed the activities of reporters with a BS-BS spacer length ranging from 4 to 14 bp. Strikingly, this spacer length affected the transcriptional activity in an apparently periodic manner for all of the four tested BSs (**Figure 3A**). The highest transcriptional activity was consistently achieved at a spacing of 5 bp, then decreased when bases were added until it reached a minimum at ∼11 bp, and then increased again with additional bases. This trend was consistently observed across the two core promoters and the three spacer sequences used in this study (**Figure 3B**), and also across all three cell types (**Figure 3C**). The apparent ∼10 bp phasing might indicate that the relative positioning of adjacent BSs on the DNA helix is a key factor for achieving high transcriptional activity.

**Figure 3:**
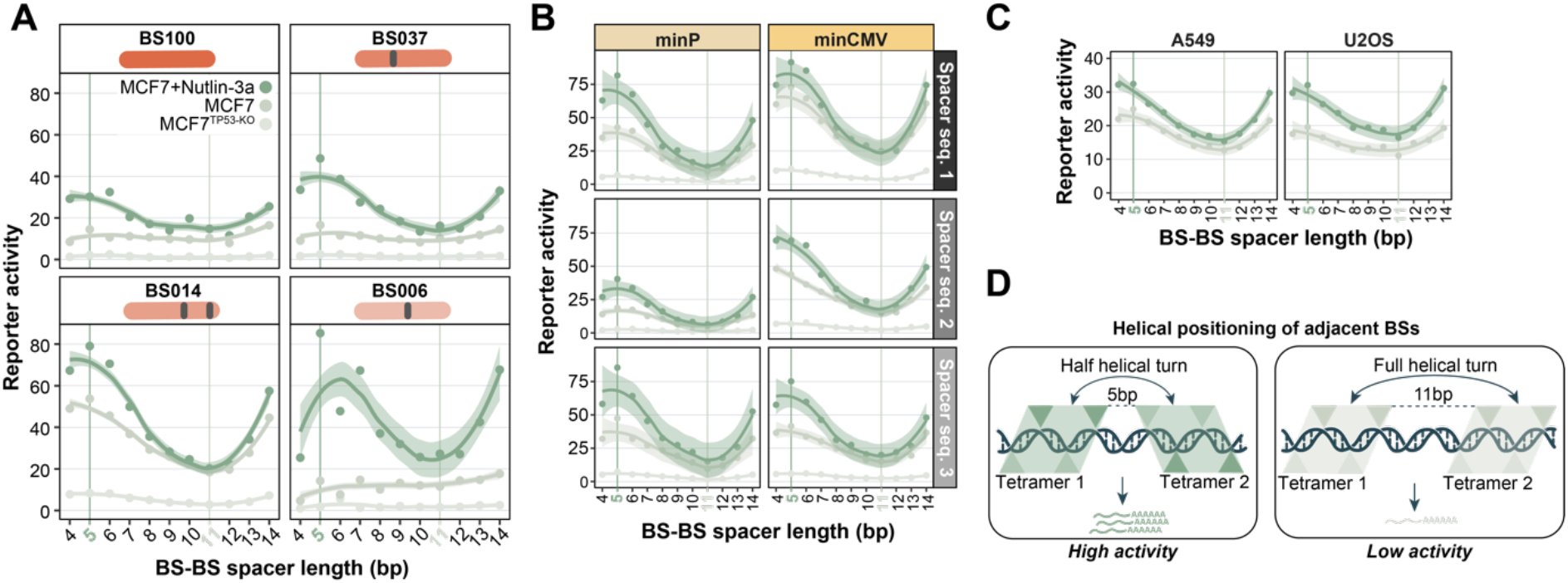
BS-BS spacer length affects transcriptional output periodically. (**A**) Activities of minCMV reporters in MCF7 cells (TP53-KO, TP53-WT and TP53-WT + Nutlin-3a) as a function of BS-BS spacer length, with four identical BSs as indicated. Activities are averaged across the three BS-BS spacer sequences. (**B**) Reporter activities (only BS014) per BS-BS spacer length, separately for the three BS-BS spacer sequences and the two core promoters. (**C**) Data shown in **B** for A549 and U2OS cells (average across BS-BS spacer sequence and core promoter). Data points in **A-C** were fitted using LOESS smoothing with a 95% confidence interval. (**D**) Model of adjacent TP53 tetramers on DNA at BS-BS spacer lengths of 5 bp (upper panel) or 11 bp (lower panel). The triangles indicate TP53 monomers.

#### Possible interpretation of BS spacing effects

A TP53 tetramer binds to its BS as a dimer-of-dimers, with two monomers on each side of the DNA double helix; the four monomers and the DNA are roughly in a single plane (33) (**Figure 3D**). At a 5 bp BS-BS spacer length, the centres of two adjacent tetrameric BSs are approximately positioned two and a half helical turns apart (because one full tetramer BS is 20 bp). Moreover, the centre of the closest monomeric BSs of two adjacent TP53 BSs are approximately one full helical turn apart. Hence, we hypothesize that this positioning on the DNA is responsible for higher transcriptional activity, either by stabilising TP53 tetramers on the DNA, or by allowing for efficient interaction with the transcription machinery, possibly increasing transcriptional burst duration or frequency (36).

### Effects of core promoter and BS-BS spacer sequences

#### Core promoter choice affects basal reporter activity and inducibility

Next, we investigated how the activity of the reporters depended on the choice of core promoter. Interestingly, the scrambled BSs were more active when coupled to the minCMV promoter (27) compared to the minimal TATA promoter (minP, derived from pGL4 (Promega, Madison, WI, USA)), demonstrating that the minCMV promoter has a higher basal activity than the minP promoter (**Figure 4A**). However, after normalisation of this basal activity, minCMV reporters with at least one functional TP53 BS were on average 1.8 and 1.5 times more active than minP reporters in MCF7 cells and MCF7 cells stimulated with Nutlin-3a, respectively (**Figure 4B**). This preferential cooperativity with minCMV was even more pronounced in A549 and U2OS cells (**Figure 4B**). Thus, despite the higher basal activity of the minCMV promoter, it still allows for higher TP53-mediated transcriptional induction. These results are contradicting a previous study showing that a minimal TATA promoter is less leaky and more inducible than a minCMV promoter (37).

**Figure 4:**
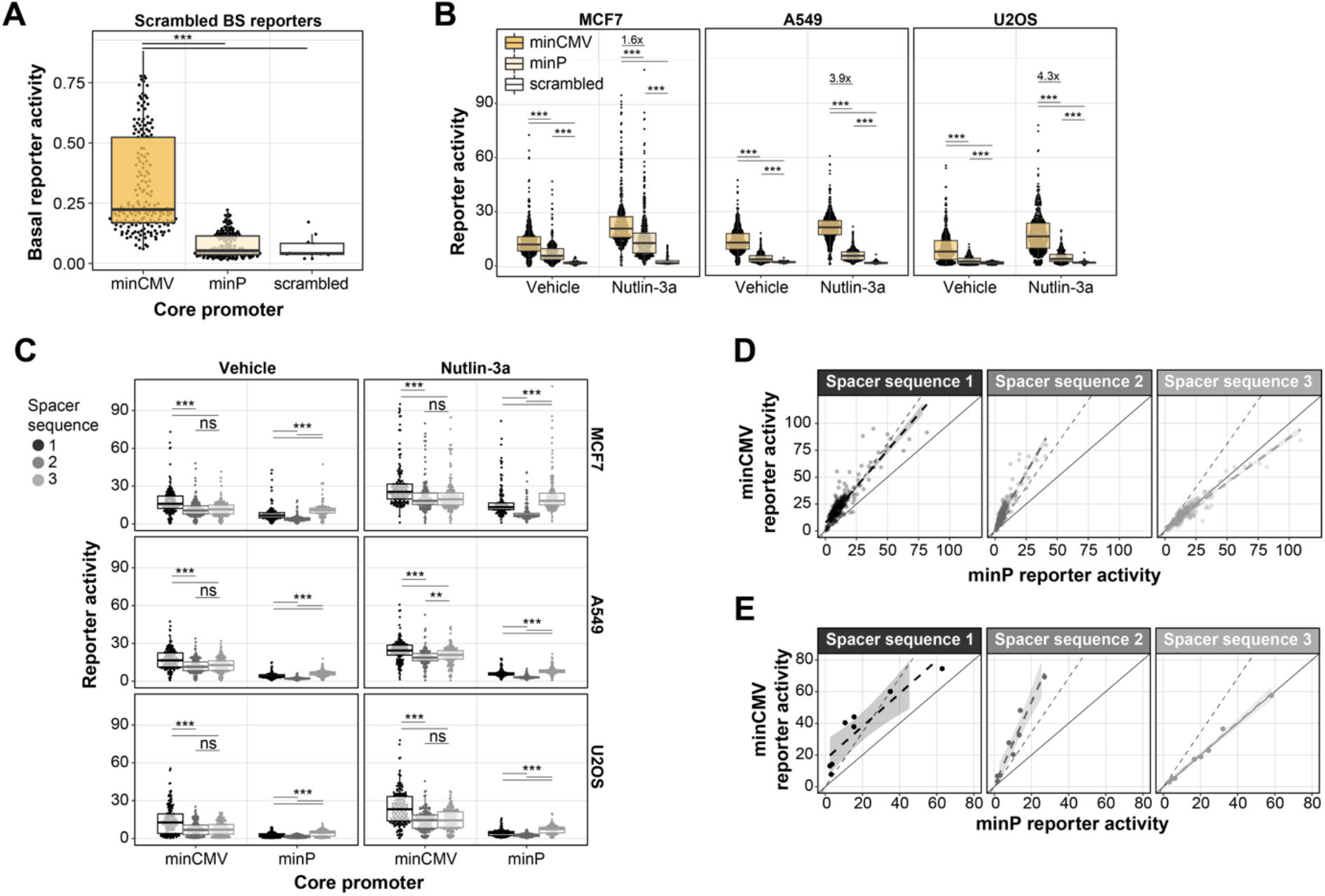
Effects of core promoter and BS-BS spacer sequences. (**A**) Activities of reporters with scrambled BSs per core promoter. Basal reporter activities are displayed and are not normalised per core promoter like in all other figure panels. (**B**) Reporter activities of all TP53 reporters per core promoter in MCF7, A549, and U2OS cells treated with vehicle or Nutlin-3a. (**C**) Reporter activities of all TP53 reporters per BS-BS spacer sequence and core promoter in MCF7, A549, and U2OS cells treated with vehicle or Nutlin-3a. (**D**) Correlation of the reporter activities between the two minimal promoters per BS-BS spacer sequence in MCF7 cells. Solid line indicates proportional relationship with slope 1, dotted line indicates the proportional relationship with the slope of the mean fold-change of minCMV over minP reporters. Data points are fitted using linear regression with a 95% confidence interval, as indicated by the grey shaded dotted line. (**E**) Same as **D** but only for reporters with a constant spacer sequence between the BSs. Groups in **A, B** and **C** were compared using Wilcoxon rank sum test, significance levels: ***: p < 0.001, **: p < 0.01, *: p < 0.05, ns: p >= 0.05.

#### BS-BS spacer sequences can modulate the core promoter activity

Even though we designed all BS-BS spacer sequences to minimise the occurrence of other TF motifs, we observed systematic differences in reporter activities between the three designed BS-BS spacer sequences (**Figure 3B, Figure 4C**). These results were concordant across all three tested cell types. When correlating the minP and minCMV reporter activities, we found that the BS-BS spacers linearly scaled the activities of all reporters in the library. Spacer *3* boosted the activity of minP reporters compared to minCMV reporters, and spacer *2* increased minCMV reporter activity compared to minP (**Figure 4D**). Importantly, the BS-BS spacer sequence showed the same phenotype when only the 10 bp spacer in front of the core promoter was varied and the spacer sequence between the BSs was kept constant (**Figure 4E**). This suggests that the 10 bp spacer sequence directly upstream of the core promoter alone may affect the reporter strength. We considered that we inadvertently created a TF motif in some of the junctions of the spacers and core promoters, but we did not find any known TF motif in these junctions.

### Explaining TP53 reporter activity using log-linear models

#### Log-linear model captures the relative importance of design features

Next, we assessed the relative contributions of the various tested features to the reporter activity. For this purpose, we constructed a log-linear model that included BS affinity, BS-BS spacer length (transformed to helical position), BS-BS spacer sequence and promoter identity, as well as an interaction term between the latter two (**Figure 5A-B**). The model was fit to the average reporter activity of the three cell types, as we wanted to explore pan-cell type effects. For simplicity we included only reporters with four identical BSs in the model. The log-linear model captured 85% and 83% of the observed variances in reporter activity in cells treated with vehicle and Nutlin-3a, respectively (**Figure 5C**, right panel). In both conditions, the helical position score had a significant positive contribution to the log-linear model, confirming that BS spacing is important for strong transcriptional activation (**Figure 5C**, left panel). As already observed in **Figure 3A-B**, the helical position has a stronger contribution to transcriptional output at high levels of TP53 (i.e., when stimulated with Nutlin-3a), suggesting that this mechanism is particularly important when multiple TP53 molecules bind at the same time to achieve high transcriptional output. Additionally, the models indicate that the lower affinity BSs BS014 and BS006 allow for high transcriptional activity (**Figure 5C**, left panel**; Figure 2A**). Finally, the models capture the importance of the BS-BS spacer sequence and promoter identity observed in **Figure 4C-E** and demonstrate the large contribution of these features compared to the helical position and the BS identity (**Figure 5C**, left panel). We note that all possible feature combinations covered by the model were also present in the reporter library. Hence, the model could not be used to identify novel feature combinations that may be highly sensitive to TP53 levels. Rather, the value of the model is that it shows that BS affinity, helical BS positioning and the BS-BS spacer sequence and core promoter identity are all important contributors to the transcriptional output; and that the relative importance of these features depends on the concentration of TP53.

**Figure 5:**
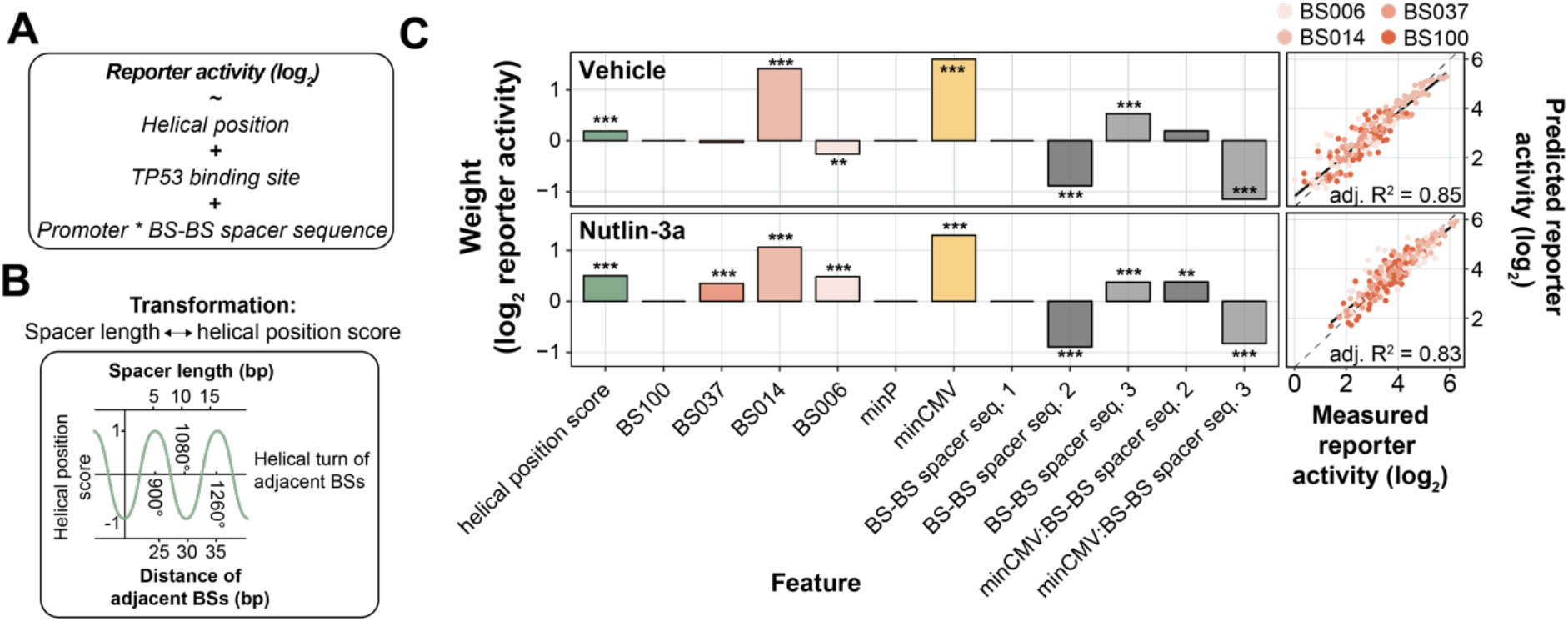
Log-linear model captures importance of design features. (**A-C**) Log-linear modelling to estimate relative contributions of BS spacing, BS-BS spacer sequences and core promoter choice. (**A**) Equation of the log-linear model used to fit the reporter activities. Reporter activities were averaged across the three probed cell types. (**B**) Explanation of the transformation of BS-BS spacer length (bp) to the helical position score. (**C**) Right-side plots: scatter plots of measured reporter activities compared to reporter activities as predicted by the log-linear model. Adjusted R^2^ is calculated from R^2^ by adjusting for the total number of model features. Left-side plots: weights calculated by the log-linear model. Asterisks indicate p-values of the individual features, significance levels: ***: p < 0.001, **: p < 0.01, *: p < 0.05, ns: p >= 0.05.

### TP53 reporters with increased TP53 sensitivity

#### Optimised reporters outperform benchmark TP53 reporters

Finally, we asked which reporters are most optimal for practical purposes, *i.e*., to measure relative TP53 activity in a standard reporter assay. Here, we considered both the signal strength of the reporters in TP53-WT cells and the dynamic range of their response to Nutlin-3a. Based on these two measures we identified two sets of TP53-sensitive reporters that can be used for different purposes. Reporter set A comprises three powerful reporters that are highly active at basal TP53 levels (49-58-fold over background on average across the three cell types) (**Figure 6A**). Under these basal conditions, reporters A1-A3 are on average 8.1 (A549), 9.2 (MCF7), and 21.7 (U2OS) times more active than the set of benchmark reporters (which produce signals 2.3-7.2-fold over background in the three cell types) (**Figure 6B**). However, set A reporters show only marginal further activation upon Nutlin-3a activation (**Figure 6A-B**), indicating that their activity approaches saturation. Therefore, we also selected three reporters (set B) that exhibit lower basal activity than set A – but still higher than all but one of the benchmark reporters – and shows an additional 4.3-5.1-fold increase in activity upon Nutlin-3a treatment (**Figure 6A**; B1-B3), and thus covers a broader dynamic range than set A. In the presence of Nutlin-3a, set B reporters are on average 2.2 (A549), 2.3 (MCF7), and 1.5 (U2OS) times more active than the benchmark reporters (**Figure 6B**). Thus, while reporters in set B are particularly useful to measure relative TP53 activities across a large dynamic range including very high TP53 levels, set A reporters may be used to sense TP53 activity at basal levels with extremely high sensitivity.

**Figure 6:**
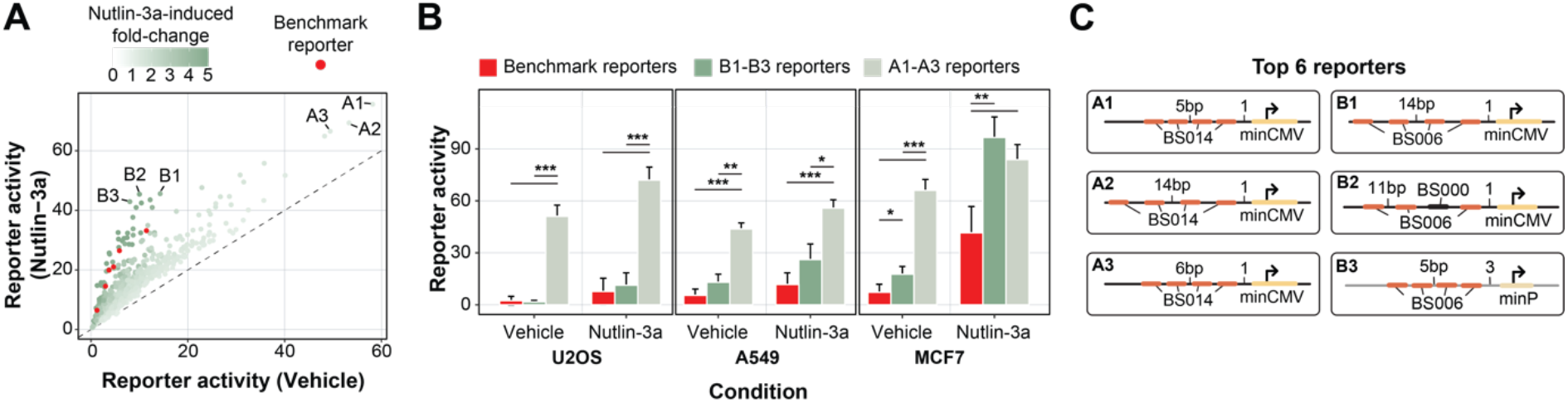
Identifying reporters with high TP53 activity and sensitivity. (**A**) Scatter plot showing the differences in reporter activity in TP53-WT versus TP53-WT stimulated with Nutlin-3a. Displayed is the mean effect across the three tested cell types. Benchmark reporters are indicated in red. The two sets of identified TP53-sensitive reporters are denoted as A1-A3 and B1-B3. (**B**) Reporter activities of the three set A (light green) and three set B (dark green) reporters versus the six benchmark reporters (red) in the different conditions. Displayed is the mean and standard deviation across the reporters. P-values were calculated using two-tailed paired Student’s t-test: significance levels: ***: p < 0.001, **: p < 0.01, *: p < 0.05. (**C**) Schematic representation of the design of the set A and set B reporters

#### Optimal reporters fit model predictions

The designs of the identified set A and set B reporters are shown in **Figure 6C**. Set A reporters all carry four copies of BS014, confirming that this BS is suitable to detect TP53 activity optimally at basal TP53 levels. Set B reporters all contain three or four copies of BS006. Furthermore, set A and set B reporters predominantly carried features with positive coefficients in the log-linear model, highlighting that the choice of BS-BS spacer lengths, BS-BS spacer sequences, and core promoter is critical to create sensitive TP53 reporters. The exception to this trend is reporter B2 with only three copies of BS006, which was not taken into consideration in the modelling.

## DISCUSSION

### Helical positioning defines transcriptional output

The results presented here provide insights into the transcriptional grammar of TP53 in the absence of other transcription factor (TF) BSs. One of the most striking findings is the effect of the relative positioning of adjacent TP53 BSs. We observed up to 4-fold differences in transcriptional activity only by changing the spacer length between adjacent BSs from 11 bp to 5 bp. Additionally, the BS-BS spacer length influenced the transcriptional activity in an apparently periodic manner, suggesting that adjacent BSs need to be positioned in a favourable helical orientation with respect to each other in order to cooperatively transcribe. Periodic helical position effects on transcriptional activity have been observed previously for other TFs (22,38,39). These insights help to understand how TP53 drives transcription from adjacently positioned TP53 BSs in a designed setting. However, it remains challenging to translate these findings to genomic TP53 response elements because of the low number of genomic TP53 response elements with adjacent BSs within the spacing range explored here, and because of the influence of confounding factors such as secondary TF BSs.

### Low-affinity BSs are strong transcriptional activators

In contrast to previous studies that explored the relation between TP53 binding affinity and transcriptional activity using *in vivo* assays like ChIP-seq (20,21,40), we focus here on affinities inferred from *in vitro* measurements. This has the advantage of studying the direct effect of binding affinity in an isolated manner without potential confounding factors like indirect TF binding, chromatin environment, locally changed TF concentrations, or interactions with other TFs. While previous studies suggested that TF-mediated transcriptional activity correlates positively with the affinity of the TFBS (24,41), our data suggest that the consensus TP53 BS (BS100) is less efficient in activating transcription than lower affinity sites (BS006, BS0014). Among all the tested reporters, optimal transcription at basal TP53 levels and strong induction upon TP53 stimulation was achieved by reporters with optimally spaced low-affinity BSs (6% relative binding affinity). Interestingly, the full-length canonical TP53 BS does not exist in the genome, and according to our calculations the majority of genomic TP53 BSs have very low binding affinity, in agreement with earlier observations (16). Therefore, full-length canonical TP53 BSs, such as BS100 or BS037 used in this study, might result in an unfavourably high binding stability (i.e., low off-rate), leading to a suboptimal transcription level and insensitivity to changes in TP53 concentration. For many TFs, low-affinity TFBSs are commonly found in the genome and are thought to be an important hallmark of transcriptional regulation (42-45). The results presented here underline the functional importance of low-affinity TFBSs.

### Alternative mechanisms influencing TP53 transcription

Besides BS affinity, it is known that additional features, like torsional flexibility of the BS, can influence transcriptional TP53 activity substantially. The consensus core sequence ‘CATG’ allows for torsional flexibility and can bind dimers stably, while ‘CAAG/CTAG’ sequences cannot (19). This feature allows ‘CATG’-containing BSs to elicit strong transcriptional activation even at low TP53 concentrations. In this study, we only included BSs with ‘CATG’ and ‘CTTG’ core sequences. Surprisingly, we found that BS014, the only BS containing a ‘CTTG’ at one half-site, was strongly active in the basal TP53-WT condition. Even though we cannot rule out that BS014 allowed binding of another unknown TF because of the higher baseline activity in the TP53-KO cells, it is possible that the ‘CTTG’ core rendered this BS sensitive to low TP53 concentrations. The exact importance of torsional flexibility and the core ‘CATG’ sequence on transcriptional activity remains to be elucidated further. Besides torsional flexibility, additional alternative mechanisms could explain the phenotypes observed in this study. For instance, changing the relative positioning of adjacent BSs might alter the binding capacity of TP53 to half-sites (46) or create additional tetrameric TP53 BSs with long internal spacer sequences (47) or partially overlapping BSs (48). Although these alternative binding modes of TP53 have been found to be less transcriptionally potent than the conventional tetrameric spacer-less binding mode (9), they were demonstrated to be functional in the human genome and thus might alter TP53 binding and transcription from the probed synthetic constructs. In future studies, it would be interesting to systematically explore the role of these alternative mechanisms.

### The effect of the number of BSs

We found that two TP53 BSs were on average 3.8 times more active compared to one BS, suggesting that TP53 molecules can cooperatively activate transcription. While a recent MPRA study suggested that transcription from two TP53 BSs is sub-additive (18), we and other TP53 reporter assay studies (9,17) observed that TP53-driven transcription can be of supra-additive nature. This finding could be important for the interpretation of the regulatory logic of genomic TP53 response elements. While the majority of TP53 target genes are already strongly activated having only one TP53 BS (20), various canonical TP53 target genes like *GADD45A, CDKN1A, BBC3, MDM2, BAX* or *DDB2* contain multiple adjacent TP53 BSs in their promoter (9,49). Our findings suggest that having a second BS drastically increases TP53 sensitivity and therefore might allow genes to be transcribed at higher rates than genes with only one BS.

### New TP53 reporters to detect TP53 activity

The two sets of three reporters that we identified are more active than the commonly used benchmark reporters at basal TP53 levels as well as high TP53 concentrations, thus detecting a larger dynamic range of TP53 activity. These reporters may thus be used as improved sensors of TP53 activity. Potentially, these reporters are also useful in detecting activity of TP53 mutants with lowered transcriptional activity, as reported before (13,14), or perhaps in the future in combination with single-cell RNA sequencing. Although in our design we aimed to minimise the probability that the reporters respond to TFs other than TP53, such imperfect specificity can never be ruled out completely, and ideally should be tested by depletion of TP53 when the reporters are used in other cell types than those reported here. The use of barcoded reporters rather than fluorescent reporters offers opportunities to multiplex a large number of reporters, *e.g*., to probe the activity of multiple TFs, in a single experiment. The combination of barcoded set A and set B reporters may also be used to achieve both high sensitivity (set A) and a high dynamic range (set B).

## DATA AVAILABILITY

Processed data and laboratory notes are available at Zenodo: https://doi.org/10.5281/zenodo.8033875. Raw sequencing data are available at Sequence Read Archive (SRA) accession PRJNA936070, https://www.ncbi.nlm.nih.gov/bioproject/PRJNA936070/.

Code is available at Zenodo: https://doi.org/10.5281/zenodo.8099025.

## ACKNOWLEDGMENTS

We thank members of our laboratories for helpful comments; the NKI Genomics and Research High-Performance Computing core facilities for technical support. We also thank Alex Fish for fruitful discussions about the binding mode of TP53 molecules, and Karin de Visser for TP53-KO cells. This work was funded by the Oncode Institute and the European Union (European Research Council Advanced Grant RE_LOCATE, 101054449). SGM is funded by EU/MUR MSCA Young Reasercher Fellowship (Next Generation EU). Views and opinions expressed are however those of the author(s) only and do not necessarily reflect those of the European Union or the European Research Council. Neither the European Union nor the granting authority can be held responsible for them. Research at the Netherlands Cancer Institute is supported by an institutional grant of the Dutch Cancer Society and of the Dutch Ministry of Health, Welfare and Sport. The Oncode Institute is partially funded by the Dutch Cancer Society.

## AUTHOR CONTRIBUTIONS

Reporter library design: CR and MT; experiments and data analysis: MT with help from SGM; manuscript writing: MT and BvS with input from CR and HJB. Project supervision: HJB and BvS.

## SUPPLEMENTARY FIGURES

**Figure S1:**
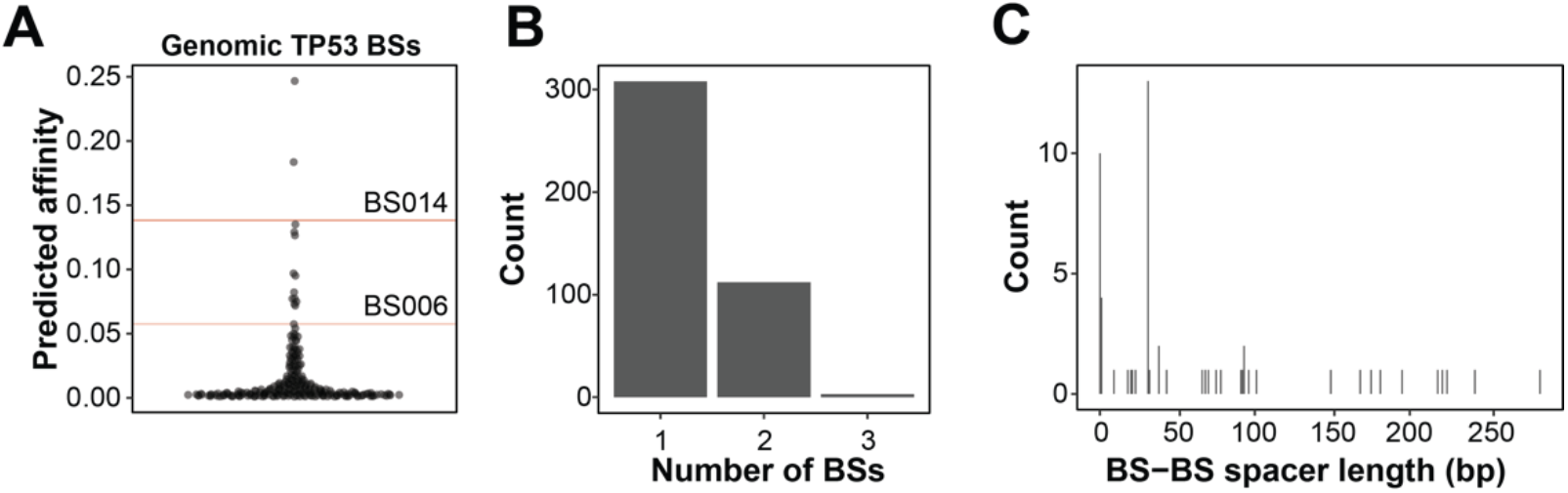
Analysis of genomic TP53 BSs. (**A**) Relative affinities compared to the consensus TP53 BS (BS100) of genomic TP53 BSs as computed by the TP53 *No Read Left Behind* model (25). Displayed are 250 BSs with the highest predicted affinities. Affinities of BS014 and BS006 are indicated by red lines. (**B**) Number of genomic TP53 response elements with one, two, or three TP53 BSs. Only BSs with a relative affinity of > 0.0001 as defined by the TP53 *No Read Left Behind* model were taken into consideration. (**C**) BS-BS spacer length distribution of all genomic response elements with two TP53 BSs.

