## Supplementary material for "Optimisation of TP53 reporters by systematic dissection of synthetic TP53 response elements": Table S1

**Supplementary Table S1:** Sequences of the primers used in the study.

| Name | Used for | Sequence (5'-3') |
|---|---|---|
| MT024 | Oligo library amplification and addition EcoRI overhang | ACCTTGGAATTCCGGAGCGAACCGAGTTAG |
| MT025 | Oligo library amplification and addition NheI overhang | CGCTTAGCTAGCCTCTTGGATGCGACGATG |
| MT165 | Gene-specific primer for cDNA synthesis | CCTCTCCGCCGCCCACCAGCTCGAACTCCAC |
| MT164 | cDNA amplification indexed primer including Illumina S1 and p7 | CAAGCAGAAGACGGCATACGAGATNNNNNNNNGTGACTGGAGTTCAGACGTGTGCTCTTCCGATCTGGTGATGCGGCACTCGATCTTCATGGC |
| MT397 | cDNA amplification indexed primer including Illumina S2 and p5 | AATGATACGGCGACCACCGAGATCTACACNNNNNNNNACACTCTTTCCCTACACGACGCTCTTCCGATCT |
